# Comparing Intra- and Inter-individual Correlational Brain Connectivity from Functional and Structural Neuroimaging Data

**DOI:** 10.1101/2024.12.03.626661

**Authors:** Xin Di, Bharat B. Biswal, Alzheimer’s Disease Neuroimaging Initiative

## Abstract

Inferring brain connectivity from inter-individual correlations has been applied across various neuroimaging modalities, including positron emission tomography (PET) and MRI. The variability underlying these inter-individual correlations is generally attributed to factors such as genetics, life experiences, and long-term influences like aging. This study leveraged two unique longitudinal datasets to examine intra-individual correlations of structural and functional brain measures across an extended time span. By focusing on intra-individual correlations, we aimed to minimize individual differences and investigate how aging and state-like effects contribute to brain connectivity patterns. Additionally, we compared intra-individual correlations with inter-individual correlations to better understand their relationship. In the first dataset, which included repeated scans from a single individual over 15 years, we found that intra-individual correlations in both regional homogeneity (ReHo) during resting-state and gray matter volumes (GMV) from structural MRI closely resembled resting-state functional connectivity. However, ReHo correlations were primarily driven by state-like variability, whereas GMV correlations were mainly influenced by aging. The second dataset, comprising multiple participants with longitudinal Fludeoxyglucose (18F) FDG-PET and MRI scans, replicated these findings. Both intra- and inter-individual correlations were strongly associated with resting-state functional connectivity, with functional measures (i.e., ReHo and FDG-PET) exhibiting greater similarity to resting-state connectivity than structural measures. This study demonstrated that controlling for various factors can enhance the interpretability of brain correlation structures. While inter- and intra-individual correlation patterns showed similarities, accounting for additional variables may improve our understanding of inter-individual connectivity measures.

## 1. Backgrounds

Studies of brain connectivity are essential for advancing our understanding of functional interactions between brain regions and the organization of the whole brain. The development of neuroimaging techniques has provided an exciting opportunity to study brain function in humans in vivo. Early research frequently employed positron emission tomography (PET) to measure glucose metabolic activity (Phelps et al., 1981) and cerebral blood flow (Fox and Raichle, 1984). These studies primarily used inter-individual correlations of PET measures to quantify brain connectivity based on glucose metabolism (Horwitz et al., 1984; Metter et al., 1984) or cerebral blood flow (Zeki et al., 1991). However, due to the nature of PET measurements, which are static, these studies were generally limited to inter-individual correlations. While they often identified statistically significant connectivity patterns, the similarities between connectivity derived from PET measures and resting-state networks identified using functional MRI (fMRI) were relatively modest (Di et al., 2017; Di and Biswal, and Alzheimer’s Disease Neu, 2012; Lizarraga et al., 2023).

Functional MRI (fMRI) has become a widely used tool for studying brain connectivity due to its superior spatial and temporal resolution (Biswal et al., 1995, 2010). Beyond capturing moment-to-moment dynamics, fMRI data can be summarized over brief periods, often during resting-state runs, to derive measures such as the amplitude of low-frequency fluctuations (ALFF) (Zang et al., 2007) and regional homogeneity (ReHo) (Zang et al., 2004). These metrics have also been applied to examine inter-individual correlations of brain (Di et al., 2024; Taylor et al., 2012; Zhang et al., 2011). Additionally, the flexibility of task designs in fMRI enables researchers to explore how task performance impacts inter-individual connectivity correlations. Studies indicate that while task conditions can induce slight changes in connectivity patterns, the overall connectivity structure tends to remain largely consistent across different tasks (Di et al., 2024).

An intriguing extension of inter-individual correlation analysis is its application to brain structural data, which tends to reflect more trait-like characteristics associated with slow and long-term effects (He et al., 2007; Mechelli et al., 2005). Mechelli and colleagues were among the first to use a seed-based approach to examine inter-individual correlations of regional brain volumes, discovering strong correlations between regions within the same functional brain systems (Mechelli et al., 2005). Building on this, He and colleagues constructed whole-brain networks based on inter-individual correlations of cortical thickness. Their findings demonstrated that these structural networks exhibit small-world properties, highlighting the efficiency and organization of the brain’s structural connectivity (He et al., 2007).

Despite its growing popularity, questions remain about whether and to what extent inter-individual correlations reflect functional connectivity, which is traditionally assessed intra-individually, typically through resting-state fMRI. One approach to validate inter-individual correlational measures is to compare their similarity to other established connectivity measures, such as intra-subject moment-to-moment functional connectivity during rest or anatomical connectivity derived from white matter tracking. When using white matter tracking from diffusion-weighted imaging (DWI) as a reference, studies have found that inter-individual correlations of structural measures show limited similarity to white matter tracts (Gong et al., 2012; Lizarraga et al., 2023). In contrast, inter-individual correlations based on functional measures of glucose metabolic activity exhibit higher similarity with white matter connectivity (Lizarraga et al., 2023). A similar pattern emerges when comparing these measures to resting-state functional connectivity. Inter-individual structural correlations display limited similarity to resting-state functional connectivity (Alexander-Bloch et al., 2013b; Di et al., 2017). However, inter-individual correlations of functional measures, such as glucose metabolic activity, show greater similiarity with resting-state connectivity patterns compared with structural measures (Di et al., 2017).

A critical question remains regarding the factors driving inter-individual variability that lead to correlations in functional or structural brain measures. Do these correlations primarily reflect individual differences shaped by genetic factors or life experiences, or do intra-individual factors also play a role? For example, inter-individual correlation analyses often include large sample sizes spanning wide age ranges, prompting the question of whether age-related effects contribute significantly to these correlations in brain structure. Exploring intra-individual correlations could provide valuable insights into the underlying causes of inter-individual variability.

In the context of functional data, such as glucose metabolism measured by FDG-PET, neural activity introduces an additional variable. This state-like factor may be influenced by participants’ mental states at the time of measurement (Di et al., 2024). Long-term brain activity, as reflected by metrics like FDG-PET or regional homogeneity (ReHo), typically persists over minutes to hours. However, it is unclear to what extent variability in this sustained activity contributes to the observed inter-individual correlations. By comparing intra-individual correlations in these slow-varying functional patterns with those from structural measures, which do not exhibit short-term fluctuations, we can gain deeper insights into the impact of state-like functional dynamics on inter-individual correlations.

In the current study, we examined correlations in brain structural and functional measures typically calculated in an inter-individual manner. We analyzed two unique datasets, allowing us to compute correlations both inter- and intra-individually, and compared the correlation structures derived from these two approaches. This comparison enabled us to estimate the contribution of intra-individual factors to the overall correlation structure. Specifically, the first dataset consists of a single individual scanned over 16 years (Duchesne et al., 2019), providing a unique opportunity to estimate gray matter and ReHo correlations intra-individually. The second dataset comes from the Alzheimer’s Disease Neuroimaging Initiative (ADNI), focusing on healthy participants with more than five longitudinal FDG-PET scans. We calculated correlations in two ways: first, by calculating correlation matrices within each participant and then averaging these matrices across participants, which minimizes individual variability and focuses on intra-individual variability; and second, by calculating inter-individual correlations at each age point and averaging these matrices across ages, which focuses exclusively on inter-individual variability while controlling for factors such as age. Lastly, we compared correlation matrices between the two datasets and investigate whether different imaging modalities have their unique correlation structures.

## 2. Materials and Methods

### 2.1 Datasets

#### 2.1.1 Simon dataset

The Simon dataset is available through the International Neuroimaging Data-Sharing Initiative (INDI) website (http://fcon_1000.projects.nitrc.org/indi/retro/SIMON.html). It includes data from a single heathy male who was scanned across 73 sessions over a 16-year period, from the age of 30 to 47. Figure 1A illustrates the distribution of these scanning sessions over time. In total, 73 MRI sessions are available, conducted using various scanners and parameters. For more details, refer to the original paper by (Duchesne et al., 2019). Our analysis focused on T1-weighted anatomical images and resting-state fMRI data, with 71 sessions providing T1-weighted images and 58 sessions containing resting-state fMRI data.

**Figure 1.**
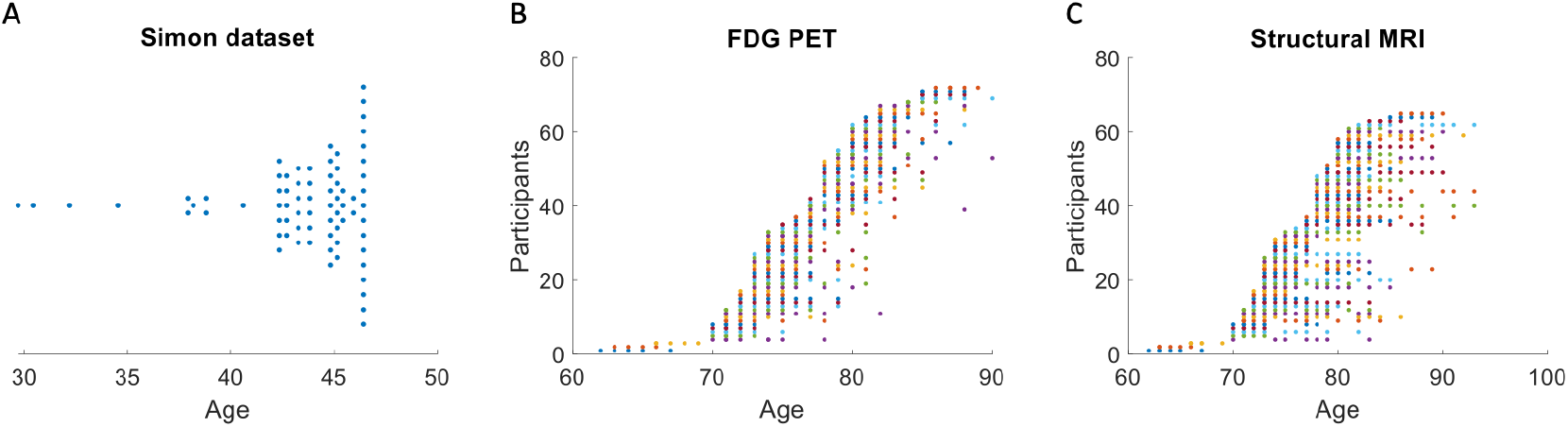
Illustration of scan sessions for the Simon dataset (A) and fludeoxyglucose-18 (FDG) positron emission tomography (PET) (B) and structural MRI (C) datasets from Alzheimer’s Disease Neuroimaging Initiative (ADNI). For the Simon dataset, a single participant was scanned for 73 sessions over 16 years. Each dot represents one session. For the ADNI dataset, each row in y axis represents one participant, where each participant was scanned for multiple sessions.

#### 2.1.2 ADNI dataset

The ADNI dataset was obtained from the project website (adni.loni.usc.edu). The ADNI was launched in 2003 as a public-private partnership, led by Principal Investigator Michael W. Weiner, MD. The primary goal of ADNI has been to test whether serial magnetic resonance imaging (MRI), positron emission tomography (PET), other biological markers, and clinical and neuropsychological assessment can be combined to measure the progression of mild cognitive impairment (MCI) and early Alzheimer’s disease (AD). For up-to-date information, see www.adni-info.org.

In this analysis, we only included data from healthy participants. All selected individuals showed no evidence of depression, mild cognitive impairment (MCI), or dementia, as indicated by Mini-Mental State Examination (MMSE) scores ranging from 24 to 30 and a Clinical Dementia Rating (CDR) of 0. We manually curated FDG-PET, MRI, and resting-state fMRI data for this study. For FDG-PET and MRI, we included participants with at least five sessions to ensure the calculation of reliable intra-individual correlations. For the resting-state fMRI data, we included only one session per individual and focused on the averaged correlation matrix across participants.

The data search began with FDG-PET, resulting in the inclusion of 72 participants (25 females) with a total of 432 PET scan sessions. The number of sessions per participant ranged from 5 to 9 (Figure 1B). The participants’ average age at the first session was 75.8 years, with a range of 62 to 86 years. For each session, either a mean image was calculated, or a single representative image was used.

From the 72 participants with qualified PET data, we identified those with at least five sessions of structural MRI scans. A total of 65 participants met this criterion, with the number of MRI sessions ranging from 5 to 13. Figure 1C provides an overview of the session count and corresponding ages for these participants.

Finally, among the 65 participants with structural MRI data, 17 individuals also had resting-state fMRI scans available. For these participants, one session per individual was included, focusing on a single resting-state fMRI session per participant. fMRI data typically lack a sufficient number of longitudinal sessions, making it impossible to conduct inter- and intra-individual correlation analyses comparable to those performed with anatomical MRI and PET data using ReHo from resting-state fMRI.

### 2.2 Data processing

Neuroimaging data processing and analysis were conducted using SPM12 (https://www.fil.ion.ucl.ac.uk/spm/) within MATLAB environment, following preprocessing and quality control procedures detailed in a prior study (Di and Biswal, 2023).

For the FDG-PET data, dynamic images (i.e., multiple images per session) were processed by realigning all images within a session to the first image, followed by generating a mean image for that session. For static PET data, which contained only a single image per session, no realignment was required. Next, the mean images (or static images) from all sessions for each participant were realigned to the image from the first session. A cross-sessional mean image was then normalized directly to the PET template in SPM, aligned to the standard Montreal Neurological Institute (MNI) space. Normalization to MNI space was performed using consistent parameters across all images. Direct normalization was chosen over MRI-mediated normalization due to the sufficient spatial resolution of PET images and the methodological advantages of direct normalization (Calhoun et al., 2017). Finally, the normalized images were spatially smoothed using an 8 mm full-width at half-maximum (FWHM) Gaussian kernel, and each image was normalized by dividing its signal by the mean signal within an intracranial volume mask.

Each anatomical image was treated as independent data, segmented into gray matter, white matter, cerebrospinal fluid, and other tissues, and normalized to standard MNI space. Spatial normalization included modulation to ensure that the resulting gray matter images reflected gray matter volume (GMV). Quality control was performed by manually inspecting the anatomical images before and after segmentation. In the Simon dataset, one session was excluded due to segmentation failure, resulting in a total of 70 sessions being included in the final analysis.

For the fMRI data, the functional images were first realigned to the first image of each session. The mean functional image was then coregistered to the corresponding anatomical image. Next, the functional images were normalized to MNI space using the deformation field maps derived from the segmentation step and spatially smoothed with an 8 mm full-width at half-maximum (FWHM) Gaussian kernel. A voxel-wise general linear model was then applied to regress out head motion parameters and white matter/CSF signals. This model included 24 regressors based on Friston’s head motion model, along with the first five principal components of signals from the white matter and cerebrospinal fluid. The residual images from this step were saved for further analysis.

For the Simon dataset, ReHo was calculated for each resting-state fMRI session using the REST toolbox (Song et al., 2011).

### 2.3 Brain parcellation

Cortical regions were defined using Schaefer’s 100-region parcellation (Schaefer et al., 2018), while subcortical regions were identified based on the Automated Anatomical Labeling (AAL) atlas. The cortical regions were further grouped into seven networks (Yeo et al., 2011) and organized by hemisphere. The subcortical regions included the bilateral hippocampus, parahippocampus, amygdala, caudate, putamen, pallidum, and thalamus (Tzourio-Mazoyer et al., 2002).

For each participant and region, voxel values were averaged to compute measures of gray matter volume (GMV), FDG-PET, and regional homogeneity (ReHo), producing a 114-dimensional vector per participant. For resting-state fMRI data, the average time series for each region of interest (ROI) was extracted, resulting in a *114* × *n* matrix, where *n* represents the number of time points, which varied across sessions and participants.

### 2.4 Calculation of intra- and inter-individual correlation matrices

In the Simon dataset, mean GMV values across 114 regions of interest (ROIs) for the 70 sessions were arranged into a 114 × 70 matrix. Pearson’s correlation coefficients were then calculated to construct a within-individual GMV correlation matrix (114 × 114). Similarly, a ReHo correlation matrix was generated using ReHo maps from 58 sessions. For each of these 58 resting-state sessions, a resting-state connectivity matrix was also computed from the fMRI time series data. Finally, the correlation matrices from all sessions were averaged to produce a mean correlation matrix.

For the ADNI dataset, correlation matrices for FDG-PET or GMV were calculated across sessions for each participant. These matrices were then averaged across participants to produce a mean correlation matrix, referred to as the intra-individual correlation matrix. To calculate inter-individual correlations, participants’ ages were controlled. Specifically, inter-individual correlations were computed at each integer age point where data from more than nine participants were available. This process was applied to participants aged between 70 and 89 years, and the resulting inter-individual correlation matrices were averaged to generate a mean correlation matrix, referred to as the inter-individual correlation matrix. Finally, for the fMRI data, resting-state functional connectivity matrices were calculated for each participant. These matrices were averaged across participants to obtain a mean connectivity matrix.

### 2.5 Fitting age models for the Simon dataset

For each region, we fitted the mean GMV or ReHo value series across sessions using cubic age models. We therefore obtained fitted age effects series and residual series across sessions. We calculated correlations of the fitted age effects and residual effects among the 114 ROIs, resulting in 114 × 114 matrices.

### 2.6 Similarities among correlation matrices

To investigate the associations among the correlation matrices, we extracted the upper diagonal of each matrix and converted it into a 6,441-dimensional vector (*114* x (*114* - *1*) / *2*). Given the potential non-Gaussian distribution of the correlation matrices, Spearman’s rank correlation coefficient (ρ) was used to quantify the relationships among the matrices.

Next, we examined the similarities between the correlation matrices from the Simon and ADNI datasets. The Simon dataset included the averaged resting-state correlation matrix, GMV correlation matrix, and ReHo correlation matrices, while the ADNI dataset contained the averaged resting-state correlation matrix, intra- and inter-individual correlations of GMV, and intra- and inter-individual correlations of PET glucose metabolic activity. To analyze these relationships, we extracted the upper diagonal of the eight matrices, forming a 6,441 × 8 matrix. We first computed Spearman’s correlation coefficients among the eight matrices, followed by a principal component analysis (PCA). Before applying PCA, the 6,441 × 8 matrix was z-transformed. We then obtained the scores of the first principal component (PC) and reconstructed them into a 114 × 114 matrix, representing a shared correlation structure across the eight matrices. Finally, we plotted the loadings of the first two PCs for the eight matrices to examine their relative positions in the low-dimensional space.

## 3. Results

### 3.1. Simon dataset

We first analyzed a multi-session dataset from a single individual spanning over 16 years. The averaged correlation matrix of resting-state time series reveals clear modular structures, evident as square-like patterns along the diagonal and additional squares representing left-right homotopic networks (Figure 2C). Subsequently, we computed intra-individual correlations for regional GMV (Figure 2A) and ReHo (Figure 2B) across all available sessions. Both matrices display square-like patterns, although their spatial configurations differ. Notably, both intra-individual correlation matrices show moderate but significant correlations with the averaged resting-state time series correlation matrix (Spearman’s correlation coefficients: *ρ*_GMV_ = *0*.*29*; *ρ*_ReHo_ = *0*.*39*).

**Figure 2.**
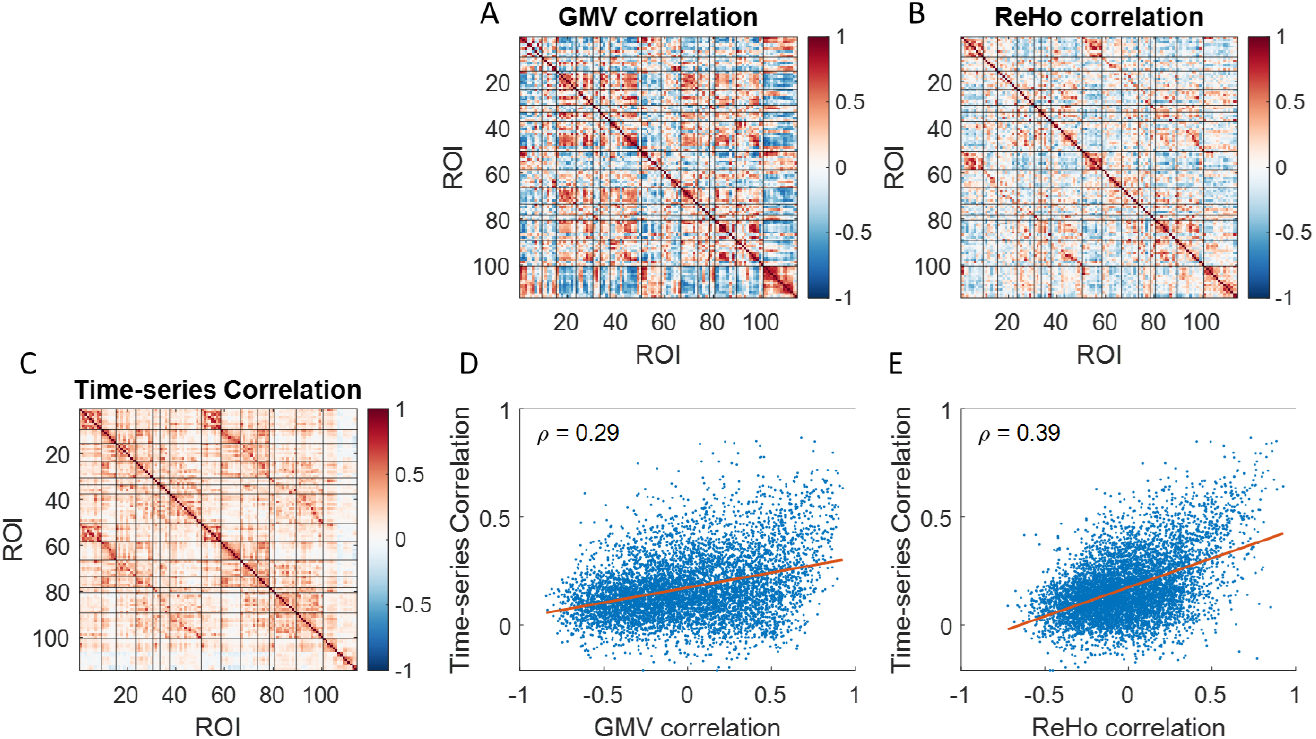
Correlations of regional brain volume (GMV) (A), regional homogeneity (ReHo) (B), and averaged correlations of resting-state time series (C) across 114 regions of interest (ROIs). Black lines delineate Yeo’s seven networks. The first 50 ROIs correspond to the left hemisphere, ROIs 51 to 100 correspond to the right hemisphere, and the remaining ROIs represent subcortical regions. D and E show the Spearman’s correlations (ρ) between the time-series correlation matrix and GMV or ReHo correlations, respectively.

To examine how long-term age effects contribute to the observed correlations, we fitted a cubic age model to the GMV time series for each ROI. Figures 3A–3C illustrate the raw time series, the fitted cubic age effects, and the residuals for all 114 ROIs, respectively. The correlation matrices for these three data types are presented in the bottom row of Figure 3. These matrices exhibited strong similarities, with Spearman’s correlation coefficients of 0.83 between the raw data and cubic age effects, 0.91 between the raw data and residuals, and 0.56 between the cubic age effects and residuals (all p < 0.001).

**Figure 3.**
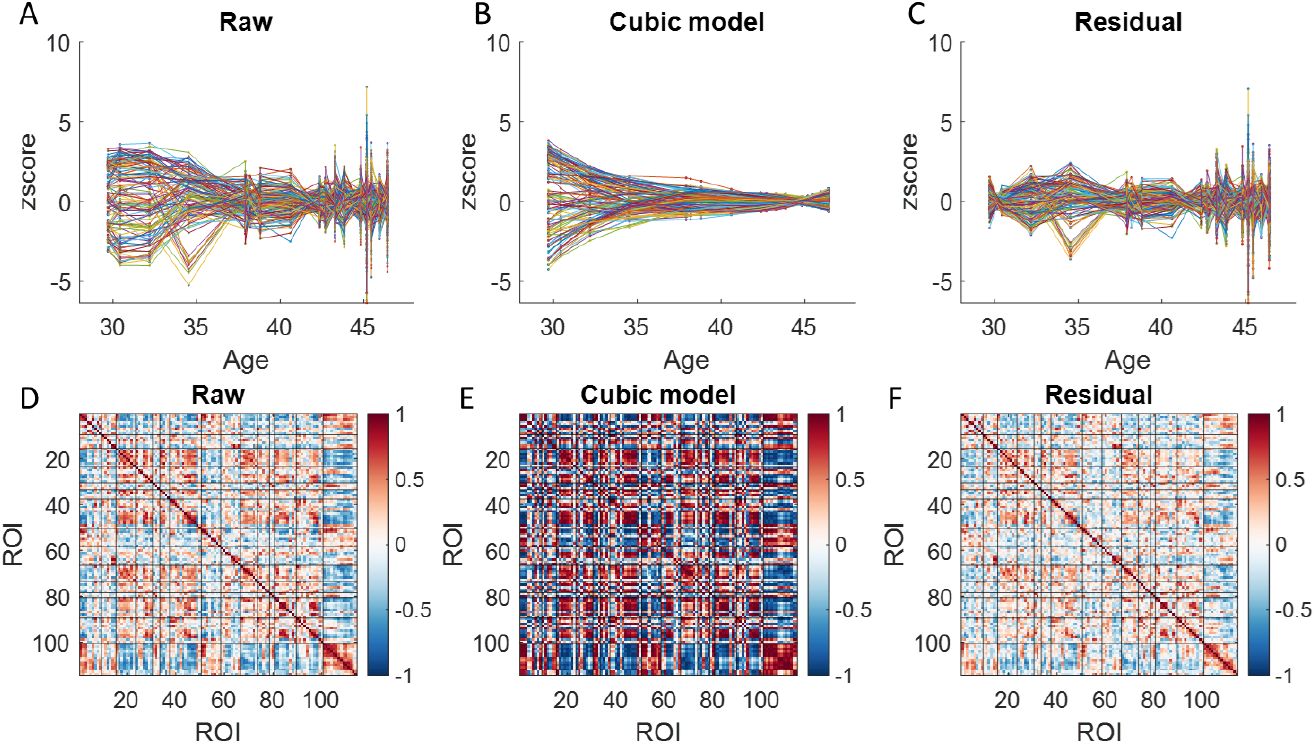
Normalized gray matter volume (GMV) for the 114 regions of interest (ROIs) (A), their fitted effects with a cubic age model (B) and the residuals (C). For illustration purpose, the time series in each ROI was converted into z values. Panels D through F, correlation matrices for the raw, fitted cubic age effects, and residuals across the 114 ROIs. Black lines delineate Yeo’s seven networks. The first 50 ROIs correspond to the left hemisphere, ROIs 51 to 100 correspond to the right hemisphere, and the remaining ROIs represent subcortical regions.

Similarly, we fitted a cubic age model to the time series of normalized ReHo values for each ROI. Figures 4A–4C display the raw time series, the fitted cubic age effects, and the residuals, respectively. The correlation matrices for these three time series are shown in the bottom row of Figure 4. All matrices were statistically significantly correlated with each other (all p < 0.001); however, the Spearman’s correlation coefficients varied considerably. The highest correlation was nearly perfect (0.99) between the raw data and residuals, while the correlations between the cubic age effects and the other two (raw data and residuals) were 0.34 and 0.21, respectively.

**Figure 4.**
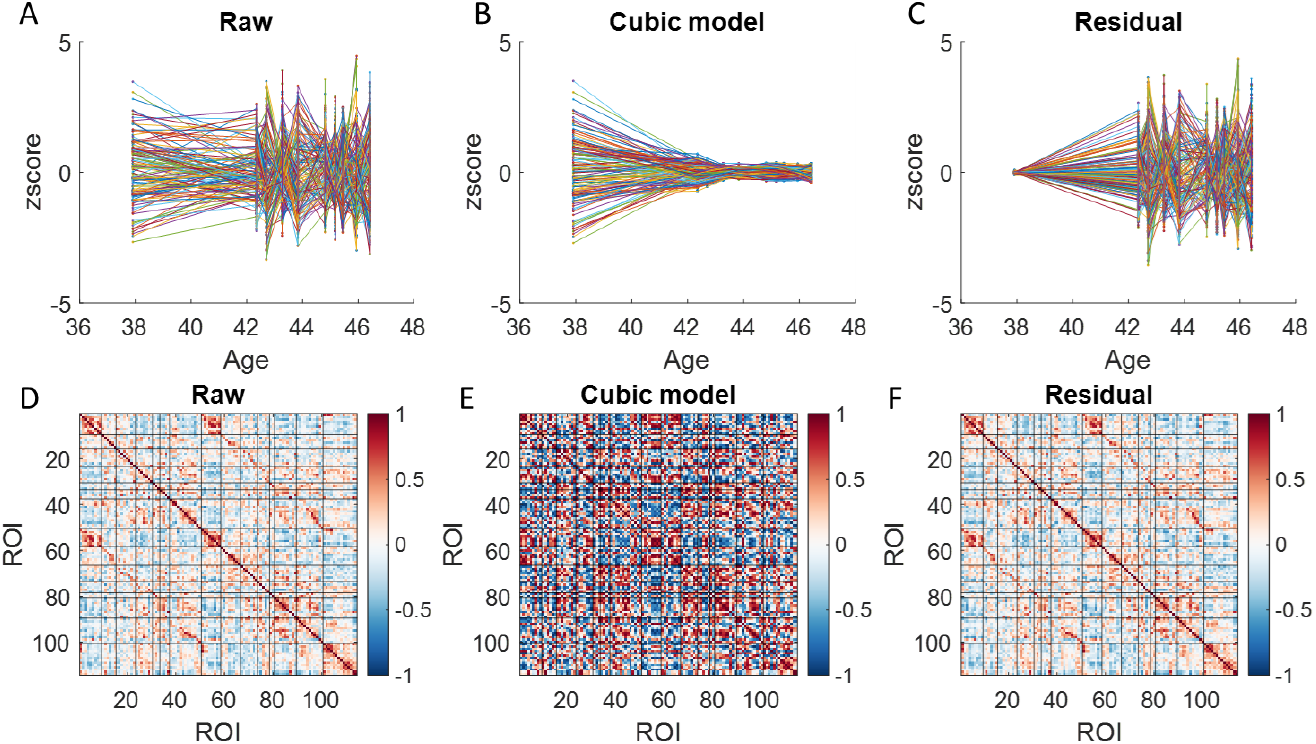
Regional Homogeneity (ReHo) for the 114 regions of interest (ROIs) (A), their fitted effects with a cubic age model (B) and the residuals (C). For illustration purpose, the time series in each ROI was converted into z values. Panels D through F, correlation matrices for the raw, fitted cubic age effects, and residuals across the 114 ROIs. Black lines delineate Yeo’s seven networks. The first 50 ROIs correspond to the left hemisphere, ROIs 51 to 100 correspond to the right hemisphere, and the remaining ROIs represent subcortical regions.

### 3.2. ADNI dataset

Next, we analyzed the ADNI dataset, where there were multiple participants, but each participant only have a few sessions. We calculated averaged resting-state time series correlation from the 17 participants (Figure 5C), which turned out to be very similar to those from the Simon dataset. We then calculated structural correlations both intra-individually (Figure 5A) and inter-individually (Figure 5B). Both matrices demonstrated functional network structures along the diagonal and between left and right corresponding regions. Moreover, both correlation matrices were correlated with resting-state time series correlation (ρ_intra_ = 0.41; ρ_inter_ = 0.37), which were slightly higher than that in the Simon dataset (0.29).

**Figure 5.**
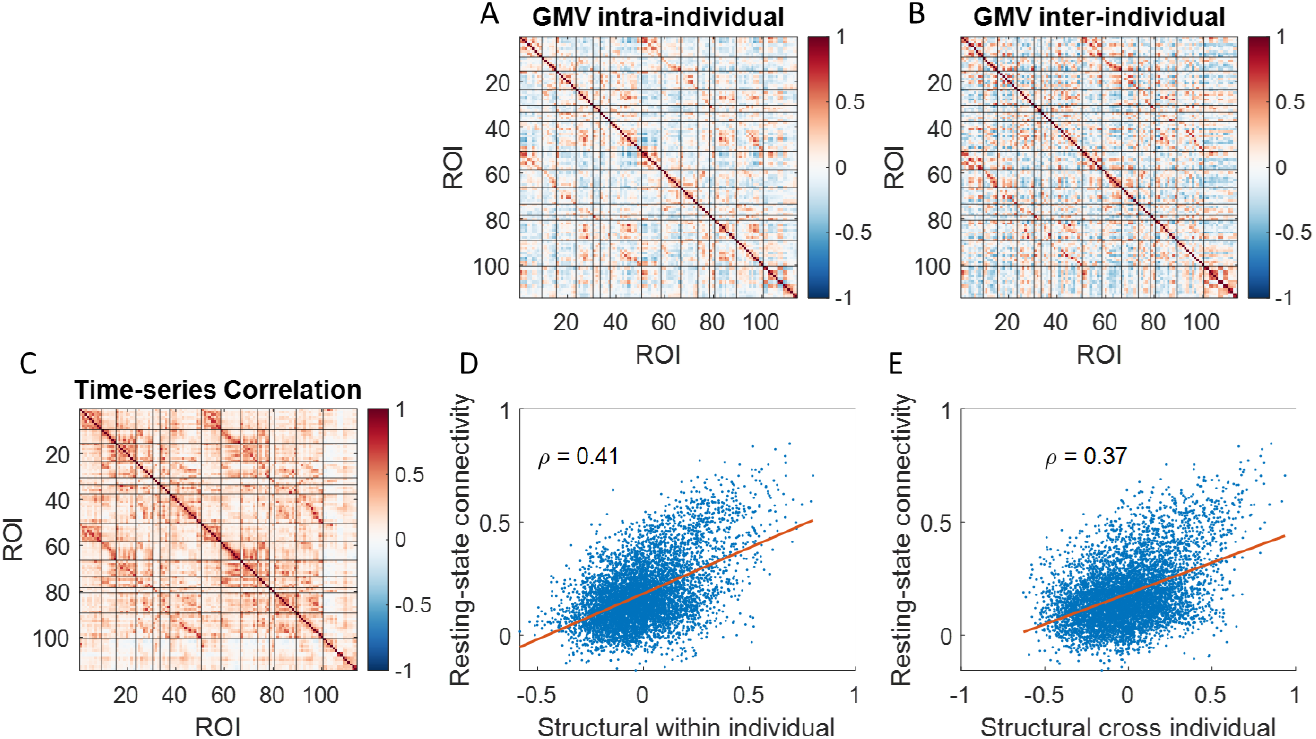
Correlations of regional brain volume (GMV) calculated intra-individually (A) and inter-individually (B), and averaged correlations of resting-state time series (C) across 114 regions o interests (ROIs). Black lines delineate Yeo’s seven networks. The first 50 ROIs correspond to the left hemisphere, ROIs 51 to 100 correspond to the right hemisphere, and the remaining ROIs represent subcortical regions. D and E show the Spearman’s correlations (ρ) between the resting-state time series correlation matrix and GMV correlation matrices, respectively.

We next calculated correlations of glucose metabolic activity intra-individually and inter-individually, and correlated the correlation matrices with resting-state time series correlation (Figure 6). The correlations matrices of glucose metabolic activity showed more obvious functional network structures than those in structural correlations. And most importantly, the both correlation matrices showed strong correlations with resting-state time series correlation (ρ_intra_ = 0.71; ρ_inter_ = 0.64).

**Figure 6.**
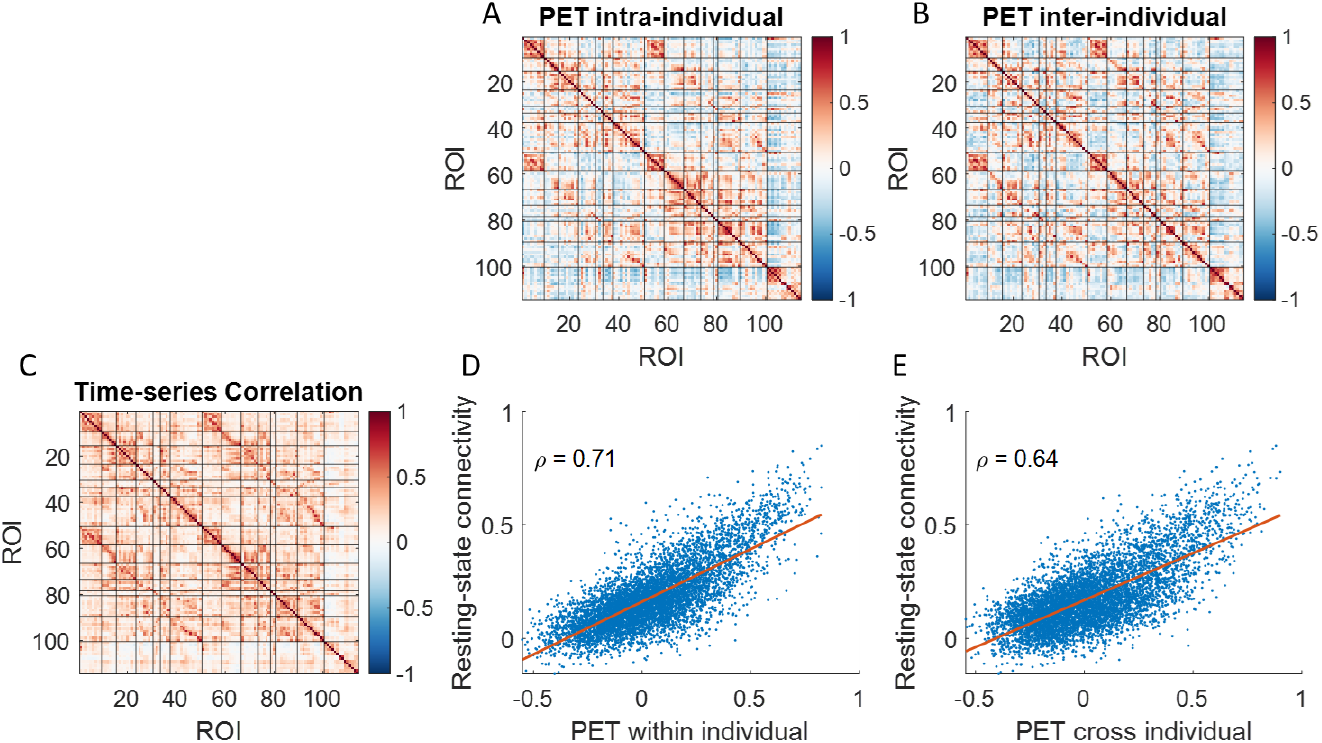
Correlations of regional brain glucose metabolism measured using positron emission tomography (PET) calculated intra-individually (A) and inter-individually (B), and averaged correlations of resting-state time series (C) across. Black lines delineate Yeo’s seven networks. The first 50 ROIs correspond to the left hemisphere, ROIs 51 to 100 correspond to the right hemisphere, and the remaining ROIs represent subcortical regions. D and E show the Spearman’s correlations (ρ) between the resting-state time series correlation matrix with the two PET correlation matrices, respectively.

### 3.3. Relationships among all the matrices

Finally, we calculated the correlations among the correlation matrices (Figure 7A). Due to the large number of ROI pairs, all correlations were statistically significant. However, it is more meaningful to focus on the relative effects of these correlations rather than their statistical significance. Here, we emphasize the correlations between the matrices of the two datasets, where the highest correlations were observed between corresponding modalities (highlighted blue rectangle and diamond markers). For instance, the strongest correlation with the mean time-series correlations in the Simon dataset was found with the time-series correlations in the ADNI dataset (ρ = 0.67). Similarly, the highest correlation with GMV correlations in the Simon dataset occurred with intra-individual GMV correlations in the ADNI dataset (ρ = 0.46), rather than inter-individual correlations. Notably, the highest correlation with ReHo correlations in the Simon dataset was observed with inter-individual PET correlations in the ADNI dataset (ρ = 0.51).

**Figure 7.**
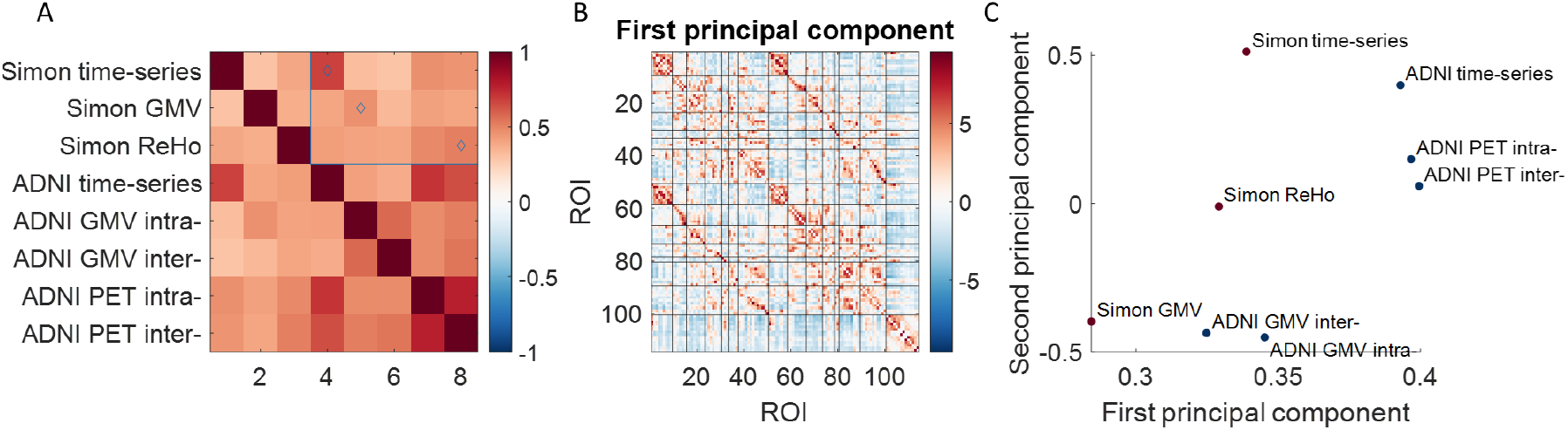
A. Spearman’s correlation matrix among the correlation matrices. The blue rectangle and diamond markers highlight the effects of interest. B. The first principal component of the correlation matrices. Black lines delineate Yeo’s seven networks. The first 50 ROIs correspond to the left hemisphere, ROIs 51 to 100 correspond to the right hemisphere, and the remaining ROIs represent subcortical regions. C. Loadings of the correlation matrices on the first and second principal components. ADNI, Alzheimer’s Disease Neuroimaging Initiative; GMV, gray matter volume; PET, positron emission tomography; ROI region of interest.

We then conducted a PCA on the eight correlation matrices, with the first principal component (PC) accounting for 59.0% of the variance (Figure 7B). Next, we visualized the loadings of the eight matrices on the first two PCs (Figure 7C). The matrices formed distinct clusters in the plot. For instance, the GMV correlation matrices were located at the bottom, while the time-series and PET correlation matrices clustered in the top-right corner.

## 4. Discussion

Using two unique datasets encompassing both intra- and inter-individual effects, the current analysis demonstrated how these effects contribute to correlation structures across brain regions. First, long-term structural brain changes revealed correlation patterns that were small but significantly associated with resting-state time-series correlations. Second, correlations in GMV were mainly driven by slow age effects, white correlations in ReHo were mainly driven by state-like brain activity. Third, long-term functional activity, as measured by ReHo or glucose metabolism, exhibited stronger correlation structures and greater alignment with resting-state time-series correlations compared to structural measures. Finally, correlation matrices from the two datasets showed greater similarity within the same modality than between different modalities, suggesting that each modality provides distinct insights into interregional relationships.

### 4.1. Structural correlations

This study analyzed a unique dataset from a single individual scanned over 16 years, revealing that intra-individual structural correlations were modestly associated with resting-state connectivity (*ρ* = 0.29), the lowest correlation observed among the analyses. These results were further validated using the ADNI dataset, which, despite fewer longitudinal data points, showed a stronger correlation between intra-individual structural correlations and resting-state connectivity (*ρ* = 0.41). These findings suggests that structural brain development and aging within an individual are partially constrained by the brain’s functional organization. Moreover, by fitting cubic age models, the fitted curve yielded correlation structures that were strongly correlated with the original GMV correlations (*ρ* = 0.83), further supported that slow aging effects could contribute to the correlation structures.

However, it is noteworthy that the variability that could not be accounted for by cubic age effects could still generate correlation matrix that were also even higher correlated with original GMV correlations (*ρ* = 0.91). Although some studies reported evidence of changes of regional brain volumes over short terms (Draganski et al., 2006, 2004), it is unlikely brain structures fluctuated with a large amount over a short term. A more plausible explanation is that the variability is due to the variability of MRI scanning settings (Di et al., 2022). For example, the homogeneity of magnetic fields may introduce spatial signal variations in a scanner, which will in turn affect spatial variations in GMV estimates. The spatial variability across scanner may introduce some correlation structures that somehow overlap with brain functional networks. This calls for proper null models for the correlations between correlation matrices (Váša and Mišić, 2022).

Notably, inter-individual structural correlations, when controlling for chronological age, showed a similar association with resting-state connectivity (*ρ* = 0.37), indicating that individual differences contribute comparably. The variability that give rise to the inter-individual correlation may be related to genetics, life experience, and plasticity (Alexander-Bloch et al., 2013a; Evans, 2013). In previous studies of structural “covariance”, these factors could not be distinguished from development and aging effects. In fact, the current findings suggest that multiple factors could give rise to the correlation structure. This is also in line with studies showing that the structural “covariance” were modulated by factors such as age (Vijayakumar et al., 2021). To enhance interpretability, researchers should consider restricting or controlling for such factors in their analyses.

### 4.2. Functional correlations

Both ReHo and FDG-PET functional correlations differed from their structural counterparts. In the Simon dataset, after removing cubic age effects, the residuals nearly perfectly mirrored the raw ReHo correlations. This indicates that ReHo correlations are primarily driven by transient, state-dependent fluctuations rather than long-term aging processes. Since ReHo summarizes neural activity during a scan session and varies with different tasks (Di et al., 2024), these fluctuations likely capture changes in mood, mental state, and daily experiences during scanning.

Additionally, correlations between ReHo or metabolic activity and resting-state connectivity were generally stronger than those for regional brain volumes. This is expected because functional measures are more directly linked to neural activity than structural ones. However, it’s important to note that neural activity measured over different temporal scales may represent distinct processes, making direct comparisons of correlation structures challenging. Previous fMRI studies have shown that connectivity patterns within a single session can vary depending on the frequency bands analyzed (Gohel and Biswal, 2015; Kajimura et al., 2023; Yuen et al., 2019). Future research should explore the characteristics of slow functional fluctuations in summary measures like ReHo and glucose metabolic activity.

### 4.3. Implications for inter-individual correlation analysis

Compared to earlier studies, the current research reports slightly higher correlations with resting-state connectivity. This improvement may stem from larger sample sizes, averaging data across multiple sessions, or advancements in the preprocessing pipeline. These findings imply that the smaller correlations observed in previous studies might partially result from noise, and that improvements in data acquisition and processing can enhance the observed similarities. Nonetheless, the analysis also highlights that each modality makes a unique contribution to the correlation structure, indicating an inherent limit to the similarities that can be achieved between different modalities.

This analysis validates the use of structural and functional brain measures to investigate interregional relationships, often referred to as functional connectivity when using functional data. Given the complex factors influencing these measures, controlling for certain variables can enhance the interpretability of correlation results. Our findings demonstrated that intra-individual correlations, which account for individual differences, tend to exhibit stronger associations with resting-state functional connectivity compared to inter-individual correlations, supporting the value of such controls. However, it is important to note that multiple measures from the same individual are not always available, and in some cases, inter-individual correlations may be the only feasible approach.

### 4.4. Limitations and future directions

One limitation of the current analysis is the narrow age range covered, making it unclear whether the findings can be generalized to other age groups. However, due to the nature of long-term longitudinal studies, obtaining suitable data across a broader age range is challenging, particularly with sufficient within-individual runs and follow-up duration. Despite this limitation, the present findings offer novel insights into the correlation structures underlying long-term effects on brain structure and function. Further studies are needed to replicate and extend these results. Another limitation is that we relied solely on resting-state functional connectivity to assess brain connectivity. Structural connectivity is typically evaluated using diffusion-weighted imaging, and previous studies have established relationships between inter-individual correlation measures and white matter connections (Gong et al., 2012; Lizarraga et al., 2023). Future studies should incorporate diffusion-based measures to create a more comprehensive picture of brain connectivity.

## 5. Conclusions

The current results to some extent validated the usage of inter-individual correlations as an estimate of brain connectivity. The results also highlighted that multiple factors could contribute to the correlation structure. Those factors may need to control or taken care of to boost interpretability of the results.

## Acknowledgement

This study was supported by grants from (US) National Institutes of Health for Xin Di (R15MH125332) and Bharat B. Biswal (5R01MH131335 and 1R01AG085665) and by (NJ) Governor’s Council for Medical Research and Treatment of Autism for Xin Di (CAUT25BRP005). Data collection and sharing for this project was funded by the Alzheimer’s Disease Neuroimaging Initiative (ADNI) (National Institutes of Health Grant U01 AG024904) and DOD ADNI (Department of Defense award number W81XWH-12-2-0012). ADNI is funded by the National Institute on Aging, the National Institute of Biomedical Imaging and Bioengineering, and through generous contributions from the following: AbbVie, Alzheimer’s Association; Alzheimer’s Drug Discovery Foundation; Araclon Biotech; BioClinica, Inc.; Biogen; Bristol-Myers Squibb Company; CereSpir, Inc.; Cogstate; Eisai Inc.; Elan Pharmaceuticals, Inc.; Eli Lilly and Company; EuroImmun; F. Hoffmann-La Roche Ltd and its affiliated company Genentech, Inc.; Fujirebio; GE Healthcare; IXICO Ltd.; Janssen Alzheimer Immunotherapy Research & Development, LLC.; Johnson & Johnson Pharmaceutical Research & Development LLC.; Lumosity; Lundbeck; Merck & Co., Inc.; Meso Scale Diagnostics, LLC.; NeuroRx Research; Neurotrack Technologies; Novartis Pharmaceuticals Corporation; Pfizer Inc.; Piramal Imaging; Servier; Takeda Pharmaceutical Company; and Transition Therapeutics. The Canadian Institutes of Health Research is providing funds to support ADNI clinical sites in Canada. Private sector contributions are facilitated by the Foundation for the National Institutes of Health (www.fnih.org). The grantee organization is the Northern California Institute for Research and Education, and the study is coordinated by the Alzheimer’s Therapeutic Research Institute at the University of Southern California. ADNI data are disseminated by the Laboratory for Neuro Imaging at the University of Southern California.

